# miRNA–mRNA Interaction Network Analysis in Alzheimer’s Disease for Biomarker Discovery

**DOI:** 10.64898/2026.01.28.702349

**Authors:** Abhishikta Ray, Komal Agarwal, Shrutika Jha, Abanindra M. Singh, Shalini Majumder, Ekarsi Lodh, Tapan Chowdhury, Alzheimer’s Disease Neuroimaging Initiative

**Affiliations:** Department of Computer Science and Engineering, Techno Main Salt Lake, Kolkata 700091, India; Department of Cancer and Genomic Sciences, College of Medicine and Health, University of Birmingham, UK

**Keywords:** Alzheimer’s disease, Transcriptomic analysis, microRNA, miRNA–mRNA regulatory network, Differential gene expression, network analysis, Neurodegeneration

## Abstract

—Alzheimer’s disease (AD) is a complex neurodegenerative disorder characterized by widespread dysregulation of gene expression and regulatory pathways. MicroRNAs (miRNAs) act as key post-transcriptional regulators by modulating messenger RNAs (mRNAs), and their disruption can influence synaptic function, neuroinflammation, and neuronal survival. In this study, we present an integrative transcriptome-driven framework to identify AD-associated miRNA–mRNA regulatory signatures and potential biomarkers. Transcriptomic and clinical data were obtained from the Alzheimer’s Disease Neuroimaging Initiative (ADNI) and the GEO dataset GSE48552. Differential expression analysis (Welch’s t-test with FDR correction) identified 148 significantly dysregulated genes (35 up-regulated, 113 down-regulated) between AD and cognitively normal controls. Experimentally validated and predicted miRNA–target interactions were integrated using miRTarBase and additional target resources, yielding 1,669,089 miRNA–gene interactions involving 3,055 unique miRNAs, with strong enrichment toward down-regulated gene targeting. Functional enrichment analysis revealed convergence of miRNA-regulated genes on synaptic signaling, neuronal communication, intracellular transport, apoptosis, oxidative stress, and PI3K–Akt/MAPK-related pathways. A bipartite miRNA–mRNA regulatory network (2,343 nodes; 14,603 edges) was constructed and analyzed using centrality metrics, highlighting key hub regulators including PBX1 and SLC7A5. Finally, supervised machine learning models trained on selected molecular features achieved strong performance, with ensemble approaches (XGBoost/LightGBM) demonstrating robust discrimination of AD from controls.

## 1 Introduction

Alzheimer’s disease (AD) is one of the most common causes of dementia and a rapidly escalating global health challenge. It is a chronic, progressive neurodegenerative disorder characterized by memory loss, cognitive impairment, and behavioral changes. According to the World Health Organization, more than 55 million individuals worldwide are currently living with dementia, with AD accounting for approximately 60-70% of these cases [1]. The irreversible nature of the disease results in severe functional decline and increased mortality, placing considerable strain on patients, caregivers, and healthcare systems. Despite extensive research, effective disease-modifying therapies remain limited due to the complex and multifactorial mechanisms underlying AD pathogenesis [2].

At the molecular level, AD is characterized by widespread dysregulation of gene expression, synaptic signaling, neuro-inflammatory processes, and mitochondrial function, indicating the involvement of multiple interacting biological pathways. MicroRNAs (miRNAs) have emerged as key post-transcriptional regulators that modulate gene expression by targeting messenger RNAs (mRNAs) [3]. These small non-coding RNAs (ncRNAs) regulate neuronal development, synaptic plasticity, immune responses, and cellular stress pathways, all of which are disrupted in AD [4]. Accumulating evidence links miRNA dysregulation to amyloid-*β*(A*β*) aggregation, τ phosphorylation, synaptic dysfunction, and neuro-inflammation [5]. Importantly, miRNAs act within complex miRNA-mRNA regulatory networks rather than as isolated regulators, motivating systems-level and network-based analytical approaches [6], [7], [8].

In this study, we present an integrative bioinformatics framework that combines differential gene expression analysis with experimentally validated miRNA-mRNA interaction data to construct AD-specific regulatory networks. Differentially expressed miRNAs and mRNAs are identified from transcriptomic data, integrated with curated interaction databases to enhance biological reliability, and analyzed using network topology measures to identify key regulatory hubs [9], [10]. Functional enrichment analysis is subsequently applied to interpret the biological relevance of the identified regulatory modules in the context of established AD-related pathways [11]. This integrative strategy emphasizes interpretability and mechanistic insight over prediction alone, aiming to uncover biologically meaningful miRNA-mRNA signatures associated with AD [12], [13].

The main contributions of this work are:

i. Identification of differentially expressed miRNAs and mRNAs associated with Alzheimer’s disease using statistically robust transcriptomic analysis.
ii. Integration of expression data with experimentally validated miRNA-mRNA interactions to construct AD-specific regulatory networks.
iii. Network-based identification of key miRNA and gene regulators acting as potential hubs in AD pathology.
iv. Functional interpretation of miRNA-regulated gene modules linked to synaptic, inflammatory, and metabolic processes.

The rest of this paper is organized as follows: Section 2 reviews related studies on transcriptomic, miRNA-based, and network-driven analyses of AD. Section 3 introduces biological preliminaries related to miRNA-mRNA regulation. Section 4 describes the datasets, preprocessing steps, differential expression analysis, target gene prediction, and network construction methodology. Section 5 presents the experimental results and network-based analyses. Finally, Section 6 concludes the paper and discusses limitations and future research directions.

## 2 Literature Review

AD has been extensively studied at the molecular level using transcriptomic, epigenomic, and regulatory network-based approaches, reflecting its complex and heterogeneous nature. The early transcriptomic studies evidenced the pervasive alterations of gene expression with implications in neuroinflammation, synaptic dysfunction, and metabolic imbalance in Alzheimer’s disease brain tissues [2]. Similarly, RNA sequencing–based investigations uniformly supported such findings and showed that many immune-related and inflammatory pathways are consistently changed throughout the disease stages, which again stresses the importance of transcriptomic analyses for understanding of AD pathogenesis [14].

Among these, miRNAs have attracted particular interest as one of the main regulators of gene expression in neurodegenerative disorders. Foundational work established that miRNAs act through post-transcriptional control of large gene networks by modulating mRNA stability and translation [3]. Systematic reviews in the context of AD reported consistent dysregulation of several miRNAs involved in amyloid precursor protein processing, tau phosphorylation, synaptic plasticity, and neuroinflammation [5].

Beyond individual miRNAs, there is growing evidence to suggest that AD is propelled by perturbations in complex miRNA-mRNA regulatory networks. Bioinformatic investigations that have jointly analyzed miRNA and mRNA expression profiles have pointed out disease-specific regulatory modules and hub genes associated with neuronal signaling and inflammatory responses [10]. Such network-centric analyses point to the limitation of singlegene approaches and show that regulatory interactions provide a far more panoramic view of the mechanisms [6]. Hence, systems biology frameworks have been accordingly proposed to unravel emergent properties of molecular networks behind neurodegeneration [7].

Network medicine further extends this view by placing AD into a systems perspective of disrupted interconnected biological systems rather than isolated molecular defects. Barabaási et al. proposed a network-based methodology to unravel modules and essential nodes of regulation responsible for bridging particular disease phenotypes [8]. The use of such knowledge regarding neurodegenerative disorders has made it feasible to pinpoint crucial molecular modules for a biomarker or a target [13].

Advantages in high-throughput transcriptomic technologies have further facilitated integrative analyses of gene regulation in AD. Best-practice guidelines in RNA-seq analysis emphasize careful normalization, differential expression testing, and rigorous statistical correction to ensure reproducibility across studies [15]. Large-scale transcriptomic studies routinely involve multiple testing, for which false discovery rate (FDR) control is commonly applied [16].

There have been an increasing number of works in this category that incorporate experimentally proven miRNA and mRNA interactions from public databases to enhance the biological plausibility of regulatory predictions. For example, the validation data added to filtering based on gene expression has enabled the predicted regulatory networks to be more biologically plausible when it comes to AD [17]. Further biological insights into the regulatory networks have also been gained by function enrichment analysis to relate the networks to the known pathways of the disease mechanisms [11].

In parallel with network-based approaches, Machine Learning (ML) and Deep Learning (DL) based models have been increasingly used within the AD setting. DL models have been distinguished for the efficiency of functionality in capturing the complex patterns of a genomic and transcriptomic nature via the acquisition of nonlinear dependencies in data with a high dimension [18]. In recent years, there have been several instances of AD classification and prediction using gene expressions with models based on ML paradigms [18], [19], [20]. Literature in recent years increasingly highlights the merging of data-driven models and biologically meaningful models. Multi-omics [21] and network studies have proved the efficiency of combining gene expressions with regulatory networks for gaining biologically actionable signs with higher efficiency in AD, compared with single-model kinds of analysis [22]. By the same mark, reviews based on miRNA signs propose about the efficiency of regulatory models in understanding about the regulation in AD signs via miRNAs [23]. Overall, prior studies highlight the value of integrating transcriptomic data with miRNA regulation and network biology to study Alzheimer’s disease. Building on this foundation, the present work applies a transcriptome-driven miRNA–mRNA network approach to identify disease-relevant regulatory signatures.

## 3 Preliminaries

### 3.1 miRNA-mRNA Interaction Axis

miRNAs are small ncRNAs produced endogenously in animals and are involved in post-transcriptional regulation of gene expression. They regulate gene expression by binding to complementary sequences in target mRNAs, leading to mRNA degradation or translational repression, thereby modulating mRNA stability and protein output in a large number of protein-coding genes in vertebrates [24]. The interaction between miRNA and mRNA has been associated with a number of human diseases, as it plays a critical role in maintaining cellular homeostasis.

Additionally, in neurodegenerative diseases, imbalanced miRNA expression manifests as misregulated control of gene expression related to neuronal function. In the context of Alzheimer’s disease, deregulation of specific miRNAs and their corresponding target gene transcripts has been consistently observed across expression profiles, indicating their involvement in disease-associated regulatory mechanisms [25]. Existing studies indicate that miRNA dysregulation in Alzheimer’s disease reflects altered regulation of established miRNAs, without evidence for disease initiation driven by novel miRNA species [26]. Constructing miRNA–mRNA regulatory networks enables systematic characterization of these interactions.

### 3.2 Degree centrality [27]

Degree centrality refers to the number of direct connections associated with a node in a network and reflects its immediate level of interaction with other nodes. For an unweighted graph *G* = (*V*, *E*), the degree centrality *d*_*i*_ of node *i* is defined as the total number of edges connected to it and is expressed as:

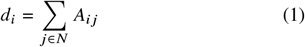

where *N* denotes the set of neighboring nodes of node *i*, and *A*_*i j*_ represents the adjacency matrix of the graph, with *A*_*ij*_ = 1 if an edge exists between nodes *i* and *j*, and *A*_*ij*_ = 0 otherwise. Nodes with higher degree centrality are considered to have greater immediate connectivity within the network.

### 3.3 Betweenness centrality [28]

Betweenness centrality quantifies the extent to which a node lies on the shortest paths between other pairs of nodes; thus, it is indicative of the potential of a node to control information flow in the network. The betweenness centrality *b*_*i*_ of node *i* is defined as:

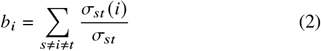

where *σ*_*st*_ represents the total number of shortest paths between nodes *s* and *t*, and *σ*_*st*_ (*i*) is the number of those paths that pass through node *i*. Nodes with high betweenness centrality are the ones that act more like critical intermediaries between different regions of the network.

### 3.4 Closeness centrality [29], [30]

Closeness centrality quantifies how efficiently a node can reach all other nodes in a network, reflecting overall accessibility within the network topology. For a graph *G* = (*V*, *E*), the closeness centrality *c*_*i*_ of a node *i* is defined as the inverse of the sum of shortest path distances from node *i* to all other nodes:

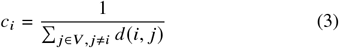

where *d* (*i, j*) denotes the shortest path distance between nodes *i* and *j*. Nodes with high closeness centrality are positioned to rapidly influence or respond to changes across the network, indicating global centrality.

### 3.5 Eigenvector centrality [31]

Eigenvector centrality assigns relative importance to the nodes based on both their direct connections and the centrality of their neighboring nodes. The eigenvector centrality *e*_*i*_ of a node *i* is defined recursively as :

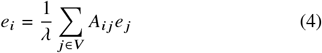

where *A*_*ij*_ represents the adjacency matrix, *e* _*j*_ is the centrality score of node *j*, and *λ* is the largest eigenvalue of the adjacency matrix. Nodes with high eigenvector centrality are considered to be influential due to their association with other highly connected and important nodes.

## 4 Methods and Materials

### 4.1 Data Resources

#### 4.1.1 Gene Expression Dataset

Transcriptomic and clinical data were obtained from the Alzheimer’s Disease Neuroimaging Initiative (ADNI) database. The expression metadata included subject identifiers (SubjectID, PTID), clinical diagnosis, demographic variables (age, gender, education), genetic risk markers (APOE4 allele count), cognitive assessment scores (MMSE), study phase, visit identifier, and RNA integrity number (RIN).

The goal of preprocessing was to create a reliable analytical dataset by removing noise, fixing inconsistencies across different data types, and keeping biologically interpretable samples that were suitable for further statistical analysis.

#### 4.1.2 miRNA-mRNA Interaction Dataset

The target genes were predicted for the differentially expressed miRNAs (DEmiRs) based on miRNA and target gene interaction data, which were extracted from miRTarBase [32], where experimental validation follows laboratory assay approaches such as reporter gene assay and expression assay, which verify miRNA regulation of target gene expression.

Datasets of target interactions were imported into a Python environment in a structured tabular format and analyzed with the `pandas` library. This study considered human-specific interaction data to maintain biological relevance with respect to AD. This is because experimentally validated interactions can avoid false-positive correlations and thus improve the quality of miRNA and mRNA regulatory networks built in this research.

### 4.2 Data Preprocessing

#### 4.2.1 Diagnostic Label Harmonization

The subjects were first categorized on the basis of diagnoses provided in the ADNI annotation. To avoid ambiguity in the diagnosis, as well as inconsistencies in the data across different time points, the diagnosis was simplified into two groups, namely, Cognitively Normal Controls (CN) and AD. To avoid overlap in molecular expression patterns, cases containing Mild Cognitive Impairment (MCI) patients in the differential expression analysis were eliminated.

#### 4.2.2 Sample Matching and Cross-Modal Integration

Clinical and transcriptomic datasets were combined using subject-level identifiers (PTID). Here, similarity-based matching was performed to fix minor formatting issues. Only samples that had a unique and clear match across datasets were kept, which ensures that each expression profile can be directly linked to a clinical record, thereby maintaining phenotype and genotype consistency.

#### 4.2.3 Probe to Gene Mapping

Probe identifiers were mapped to gene symbols using the *Symbol* annotation field. Probes associated with multiple gene symbols (e.g., “CTAGE6 || CTAGE15”) were retained to avoid information loss, while probes lacking valid gene annotations were excluded from further analysis.

Expression values corresponding to probes mapped to the same gene symbol were aggregated to obtain a single representative gene-level expression value. The resulting gene expression matrix was subsequently used for differential expression analysis. Differentially expressed genes (DEGs) were intersected with miRNA target genes to identify disease-relevant miRNA–mRNA regulatory relationships, followed by FEA to characterize associated biological processes and pathways.

#### 4.2.4 Expression Data Quality Control

For each sample *S*, the total expression was computed as:

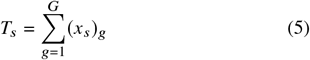

where (*x*_*s*_) *g* denotes the expression level of gene *g* in sample *s*, and *G* represents the total number of genes.

Samples with total expression values exceeding three standard deviations from the cohort mean were flagged as potential outliers. The gene-level filtering was applied as follows:

- Genes with missing expression values in more than 20% of samples were excluded.
- Low-expression genes were removed by requiring expression levels above a predefined threshold in at least 10% of the samples.

#### 4.2.5 Exploratory Data Analysis

Principal Component Analysis (PCA) was performed on standardized expression data to assess global variance structure and identify potential batch effects. PCA also served as a qualitative validation step, verifying whether diagnostic groups exhibit separable transcriptional patterns.

#### 4.2.6 Preprocessing and Standardization of Target Gene Lists

Gene symbols from the database were standardized to a uniform format using `pandas` string manipulation functions. Duplicate miRNA–gene pairs were removed to avoid redundancy. This preprocessing step ensured consistency across databases prior to integrative analysis.

### 4.3 Differential Expression Analysis

Let *x* ∈ ℝ^*N*×*G*^ denote the filtered, log_2_-transformed gene expression matrix, where *N* represents the number of samples and *G*denotes the number of genes. Each sample belongs to one of two diagnostic groups: Normal (CN) or diseased (AD). Differential expression analysis was formulated as a gene-wise statistical comparison problem aimed at identifying genes whose expression distributions differ significantly between these two diagnostic groups.

#### 4.3.1 Log_2_ Fold Change Estimation

For each gene g, mean log_2_ expression values were computed independently for AD and control samples:

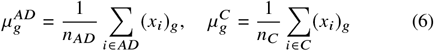

The log_2_ fold change (Log2FC) was defined as

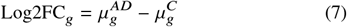

Log_2_ transformation improves variance stabilization and facilitates symmetric interpretation of up- and down-regulation. The corresponding linear fold change was derived as:

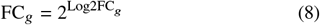

#### 4.3.2 Hypothesis Testing and Multiple Testing Correction

For each gene g, differential expression was assessed using Welch’s two-sample *t* -test applied to log_2_-transformed expression values. The hypotheses were defined as:

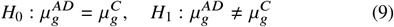

Given the large number of simultaneous gene-wise tests, raw *p*-values were adjusted to control the false discovery rate (FDR). The Benjamini–Hochberg procedure [33] was employed:

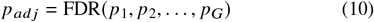

#### 4.3.3 Significance Criteria and Gene Classification

Genes were classified as differentially expressed if they satisfied both statistical and biological significance criteria:

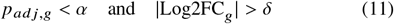

where α denotes the adjusted significance threshold and *δ* represents the minimum biologically meaningful effect size.Genes satisfying an adjusted *p*-value < 0.18 and an absolute log_2_ fold change greater than 0.25 were considered differentially expressed. Genes with positive Log2FC values were classified as up-regulated in AD, while negative values indicated down-regulation.

The differential expression workflow employed in this study is summarized in Algorithm 1.

##### Algorithm 1

Differential Expression Analysis (AD vs Control)

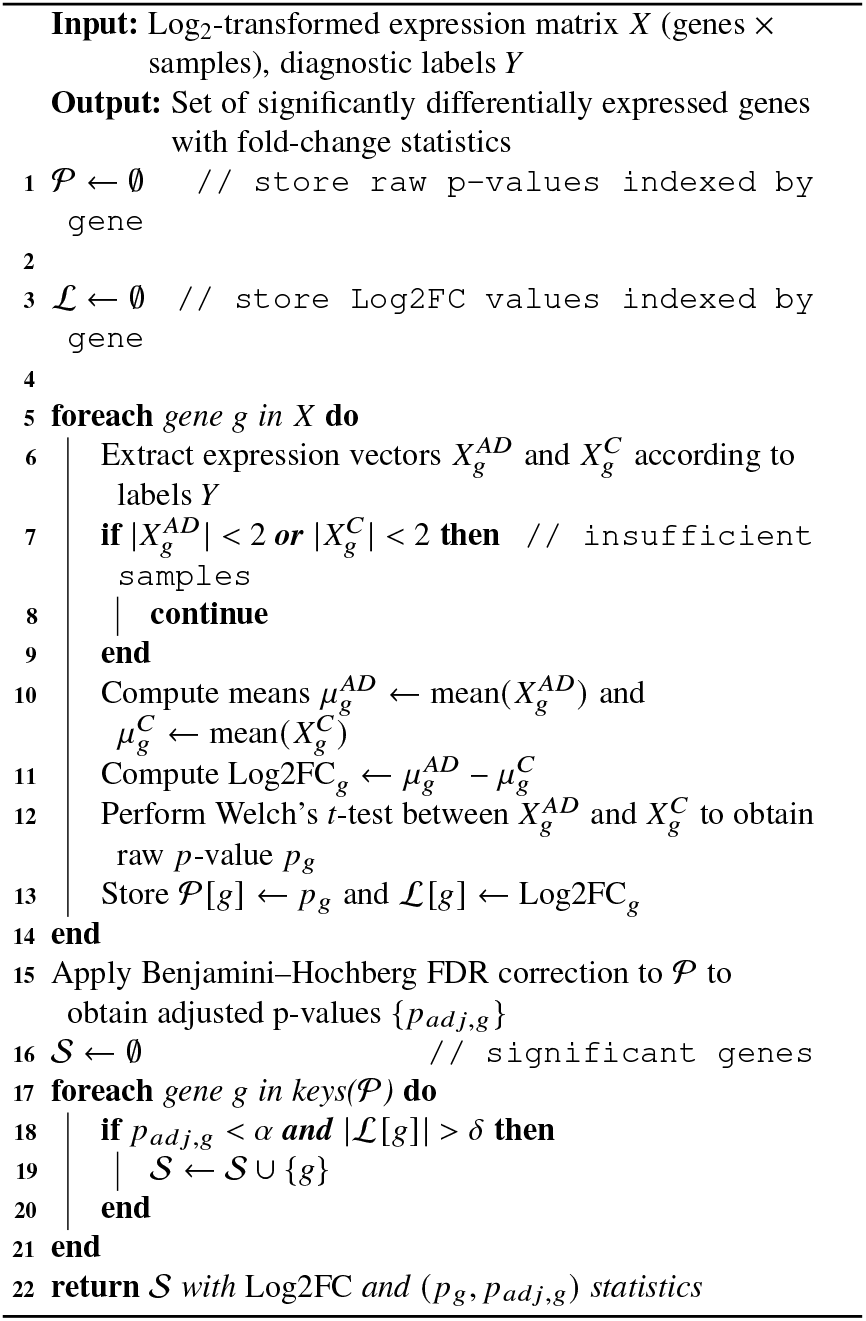

Algorithm 1 performs gene-level differential expression analysis on log_2_-transformed expression data to compare AD and CN samples, followed by a statistical assessment using Welch’s *t*-test with FDR correction. Genes meeting both adjusted significance and fold-change thresholds are retained as differentially expressed.

### 4.4 Target Gene Prediction

To identify disease-relevant regulatory interactions, predicted miRNA targets were intersected with the overall gene set. Genes present in both the overlapping gene list and differentially expressed genes (DEGs) obtained from Section 4.3 set were considered putative functional targets of dysregulated miRNAs in AD.

#### Algorithm 2

miRNA Target Gene Prediction and DEG Overlap Analysis

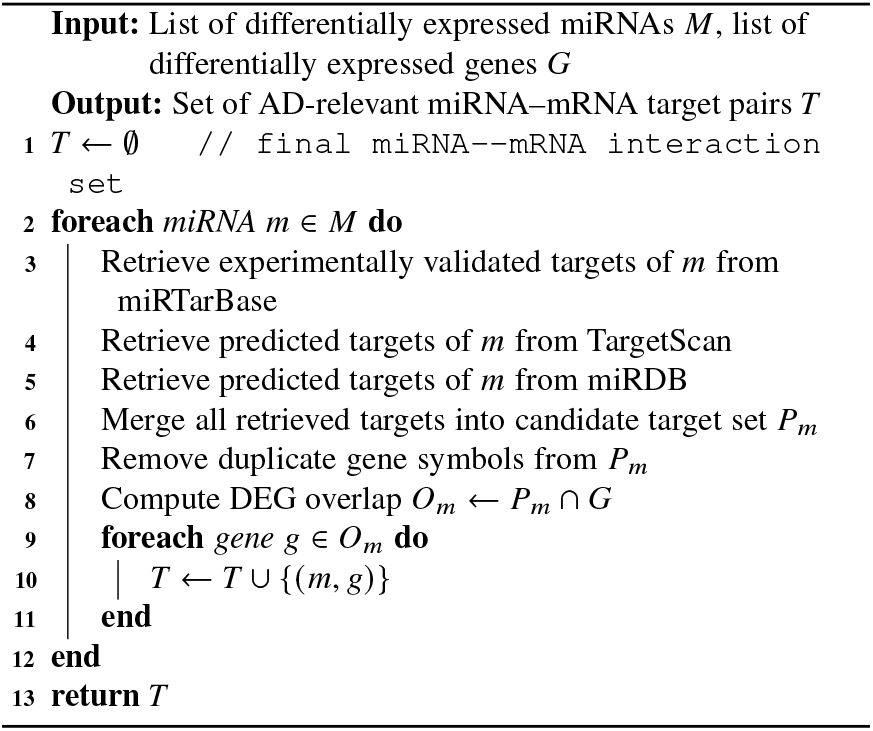

Algorithm 2 combines miRNAs that are differentially expressed with DEGs to identify AD-relevant miRNA-mRNA regulatory pairs. Validated and predicted miRNA targets are gathered from various sources and intersected with the set of differentially expressed genes. Overlaps between miRNAs and genes are kept. The overlaps represent miRNA-gene pairs that can be associated with AD.

### 4.5 Functional Enrichment Analysis

#### 4.5.1 Gene Ontology (GO) Enrichment

FEA of the overlapping target genes was performed using the `gprofiler-official` [11] Python package, which interfaces with the g:Profiler web service. GO enrichment was assessed across:

- Biological Process (BP),
- Molecular Function (MF),
- Cellular Component (CC).

The enrichment analysis employed a hypergeometric test with Benjamini–Hochberg FDR correction [33]. GO terms with adjusted *p*-values less than 0.05 were considered statistically significant.

#### 4.5.2 KEGG Pathway Enrichment Analysis

In addition to GO analysis, Kyoto Encyclopedia of Genes and Genomes (KEGG) pathway enrichment was conducted using g:Profiler. This analysis enabled the identification of signaling pathways significantly associated with the miRNA-regulated DEG set.

Enriched pathways were filtered and ranked based on adjusted significance values and gene counts.

Algorithm 3 applies FEA to the set of overlapping miRNA target genes to highlight biologically relevant GO terms and KEGG pathways. Enrichment significance is calculated using g:Profiler with Benjamini–Hochberg FDR correction. Statistically significant functional categories are retained and ranked so as to facilitate biological interpretation of AD–associated regulatory mechanisms.

##### Algorithm 3

GO and KEGG Functional Enrichment Analysis (FEA)

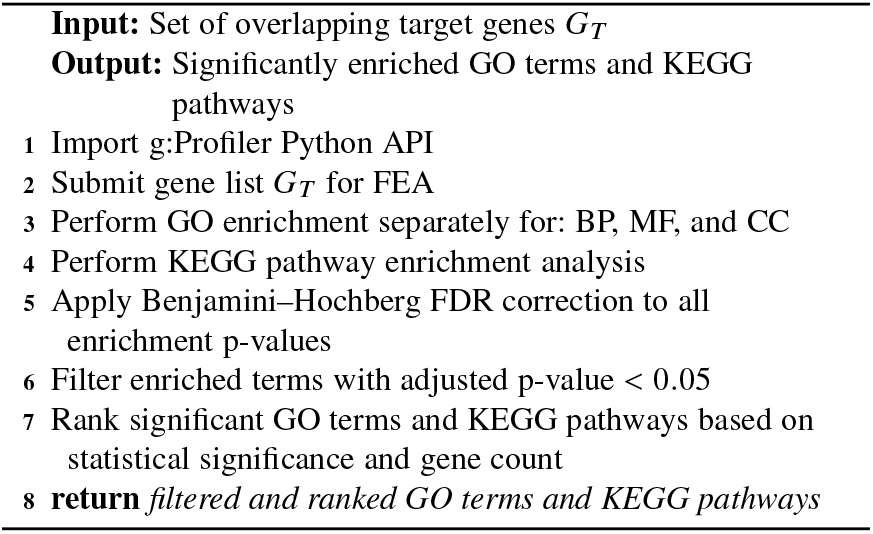

### 4.6 miRNA–mRNA Network Construction

#### 4.6.1 Network Assembly

A miRNA–mRNA regulatory network was constructed using the `networkx` [34] Python library. In this network:

- Nodes represent miRNAs or target genes,
- Edges represent validated or high-confidence regulatory interactions.

The edge list was generated programmatically from the filtered miRNA–target interaction dataset, ensuring reproducibility and scalability.

#### 4.6.2 Topological Analysis of Regulatory Networks

To identify key regulatory elements, network topological properties were computed using built-in `networkx` functions. Degree, betweenness, closeness, and eigenvector centrality were calculated for all nodes.

Nodes with high centrality values were classified as hub regulators, indicating their potential importance in coordinating transcriptional dysregulation in AD.

#### 4.6.3 Centrality-Based Backbone Extraction

To isolate the most influential regulatory structures, backbone sub-networks were extracted independently for each centrality metric by retaining top-ranked nodes and their immediate neighborhoods. These reduced representations preserve key regulatory relationships while minimizing network complexity.

#### 4.6.4 Network Visualization

The constructed network was visualized using force-directed layouts to highlight highly connected miRNAs and genes, which in turn supports the identification of central regulators and densely connected functional modules.

In Algorithm 4, a miRNA-mRNA regulatory network is built by creating nodes and edges for interactions in a graph. The topology of the graph can be measured by degree and betweenness, which reveal the importance of each node. The highly ranked nodes are also recognized as potential regulatory hubs in AD.

### 4.7 Biomarker Identification

#### 4.7.1 Proposed Feature Selection Strategy

Based on the results from Sections 4.3–4.5, candidate biomarkers were defined as:

##### Algorithm 4

miRNA-mRNA Regulatory Network Construction and Analysis

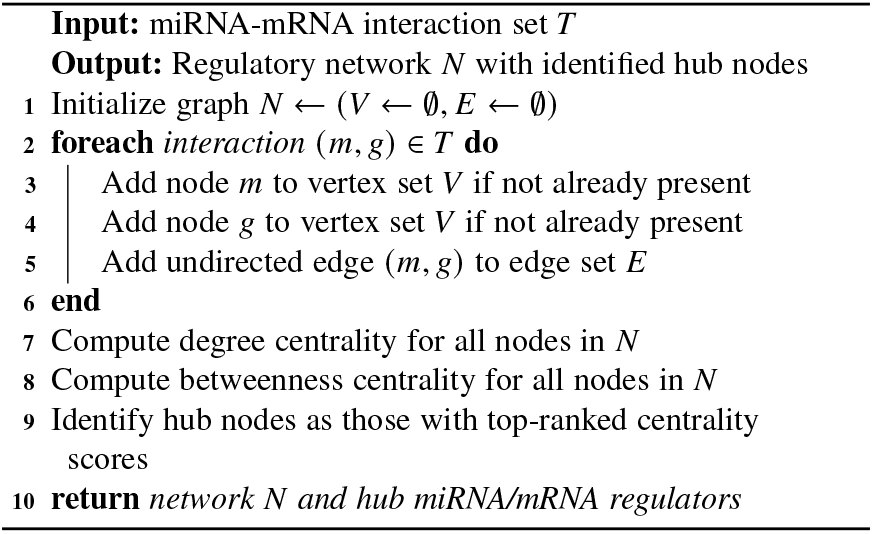

- Hub miRNAs identified from network analysis,
- Overlapping target genes with significant enrichment and high centrality.

These features were intended for use in supervised machine learning classification models.

#### 4.7.2 Classification Models

The machine learning framework was implemented using the `scikit-learn` [35] ecosystem, focusing on ensemble-based learning for the classification of AD. A **Light Gradient Boosting Machine (LightGBM)** classifier was used as it is suitable for high-dimensional transcriptomic data and can model nonlinear feature interactions explicitly. To handle class imbalance, the Synthetic Minority Over-sampling Technique (SMOTE) was applied to the training data. Model performance was evaluated with stratified *k*-fold cross-validation, and main hyperparameters were optimized through systematic tuning. Multiple models were combined using **weighted voting** and **weighted ensembling**, where weights were assigned according to the performance of each individual classifier to enhance robustness and reduce model variance while improving generalization.

#### 4.7.3 Model Evaluation Metrics

The performance of the classification models was assessed by different complementary metrics to ensure a balanced assessment of predictive accuracy and class discrimination. Overall classification accuracy was used to quantify the proportion of correctly classified samples, while precision and recall were employed to assess the reliability of positive predictions and the ability to reveal AD cases, respectively. The F1-score was computed as the harmonic mean of precision and recall to account for potential class imbalance.

Further, ROC-AUC was used as a metric to assess the discriminative capability of the model at different decision thresholds. Together, these metrics offer a solid and interpretable basis on which to evaluate the performance of this model in distinguishing AD samples from CN controls.

This algorithm is used to train supervised ML classifiers using chosen gene or miRNA features to differentiate between AD and CN samples. Models are evaluated on held-out test data using standard performance metrics, including accuracy, ROC–AUC, and F1-score. The resulting models and metrics are used to assess the predictive utility of transcriptomic biomarkers.

##### Algorithm 5

Machine Learning Framework for Biomarker Identification

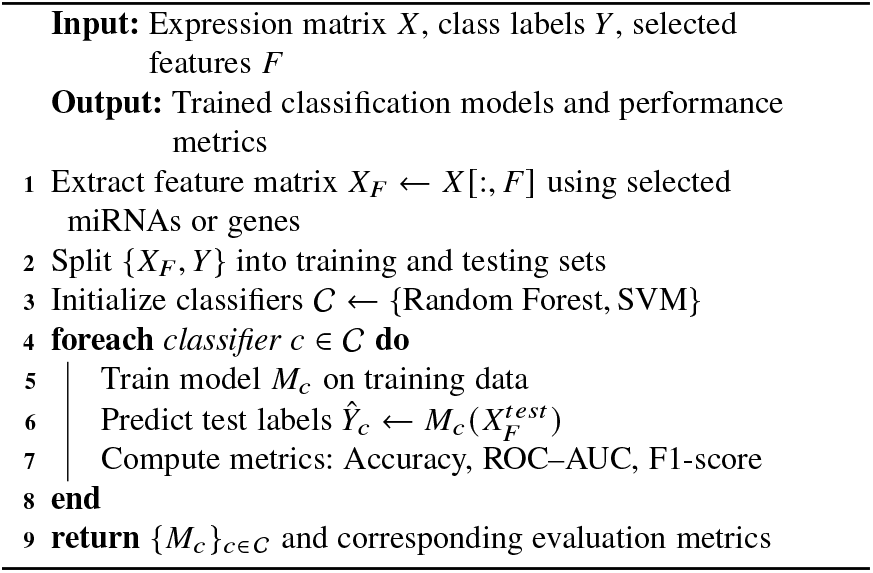

## 5 Result and Discussion

### 5.1 Differential Expression Analysis

#### 5.1.1 Integration of Gene Symbols

Probe identifiers were mapped to gene symbols, and this resulted in 48,157 valid probe-to-gene symbol mappings, corresponding to 97.5% of the original probe sets, indicating excellent annotation coverage and data integrity. Following annotation, the standardized gene symbols were used for the DEGx analysis. This step ensured that each DEG could be represented at the gene level rather than at the probe level, facilitating biologically interpretable downstream analyses.

#### 5.1.2 Differentially Expressed Genes (DEGs)

A total of 148 genes were identified as significantly differentially expressed, of which 35 were up-regulated, and 113 were down-regulated in AD samples relative to controls.

Fig. 1 illustrates an overview of the differential expression analysis between diseased and control samples using a volcano plot (left) and an MA plot (right). In the volcano plot, significantly up-regulated genes are shown in red and down-regulated genes in blue, based on adjusted *p*-value and log_2_ fold-change thresholds, while non-significant genes are shown in gray. The MA plot displays the relationship between mean expression and log_2_ fold change, highlighting systematic expression shifts and confirming the symmetry and reliability of the differential expression analysis. Table 1 provides a quantitative overview of the differential expression results. The strong predominance of down-regulated genes suggests a widespread suppression of transcriptional programs associated with disease pathology, consistent with neuronal dysfunction and synaptic loss reported in neurodegenerative disorders.

**Table 1:** Summary of Differential Expression Analysis Parameters and Results (Diseased vs Control)

| Analysis Parameter | Value |
| --- | --- |
| Total probes analyzed | 49,386 |
| Total genes analyzed | 20,092 |
| Total samples | 302 |
| Control samples | 259 |
| AD samples | 43 |
| Statistical method | unpaired *t*-test |
| *p*-value threshold | 0.18 |
| Log<sub>2</sub>FC threshold | 0.25 |
| Total significant genes | 148 |
| Up-regulated in AD | 35 |
| Down-regulated in AD | 113 |
| Percentage significant (%) | 0.74 |
| Maximum absolute log<sub>2</sub>FC | 0.9231 |
| Maximum linear fold change | 1.90-fold |
| Median absolute log<sub>2</sub>FC | 0.0321 |

**Fig. 1:**
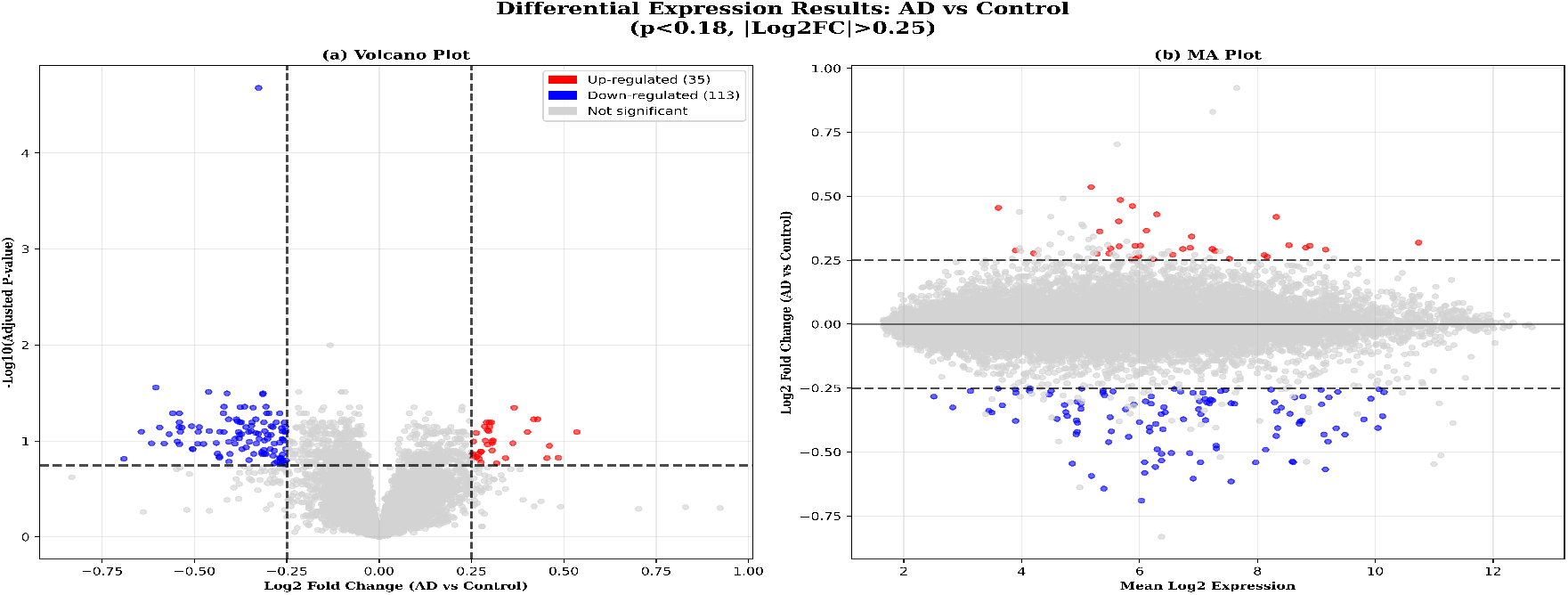
Volcano plot (left) and MA plot (right) illustrating differential expression patterns between AD and control samples. Red points denote significantly up-regulated genes, blue points represent significantly down-regulated genes, and gray points indicate non-significant genes based on adjusted *p* < 0.1 and |log_2_ |FC> 0.3. Vertical and horizontal dashed lines indicate fold change and significance thresholds, respectively.

#### 5.1.3 Statistical Significance of DEGs

Fig. 2 provides an in-depth statistical characterization of the identified DEGs. The *p*-value distribution exhibits enrichment of low *p*-values below the 0.18 threshold, indicating the presence of a genuine signal rather than a uniform null distribution.

**Fig. 2:**
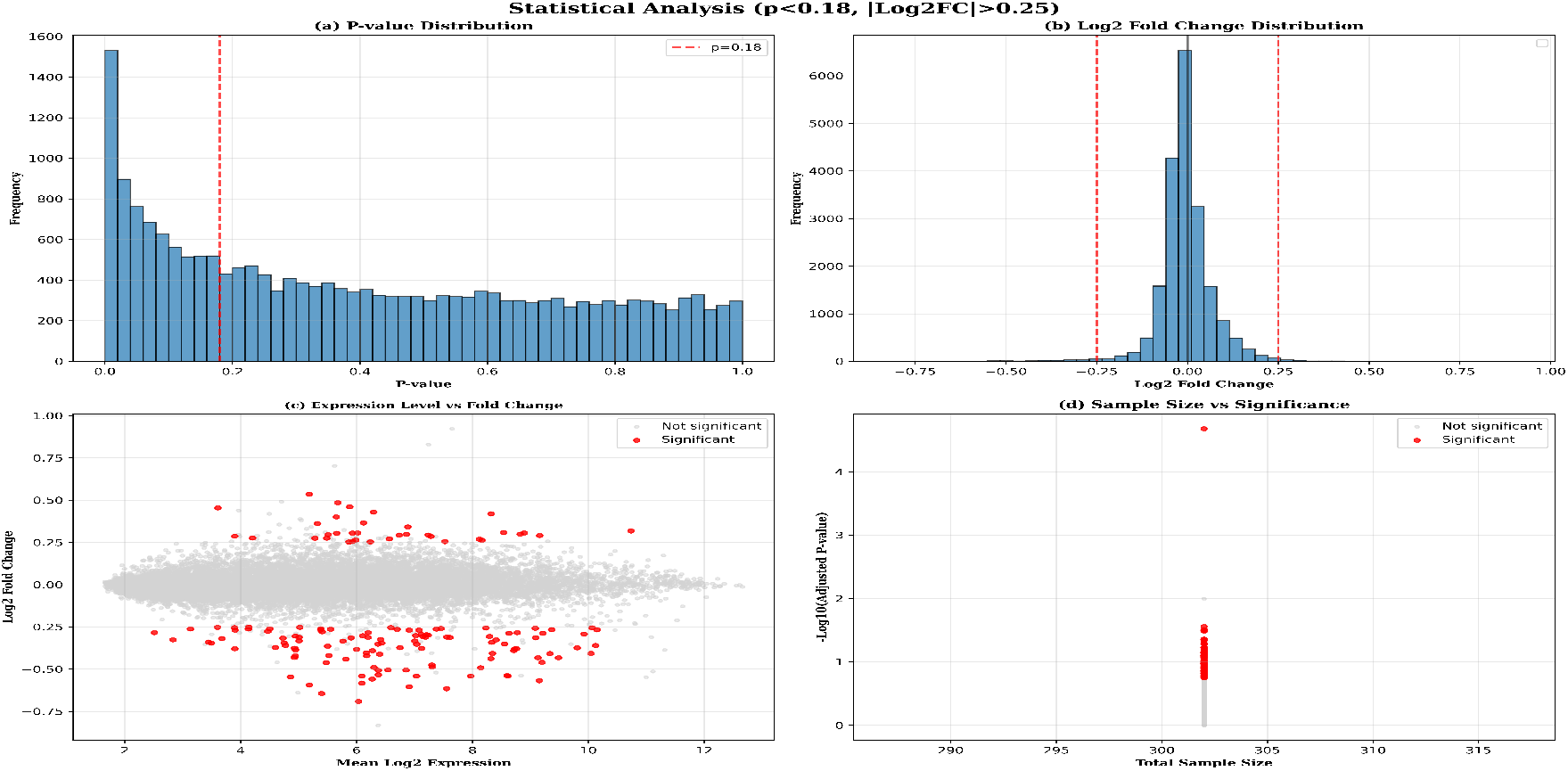
Statistical distribution plots summarizing differential expression analysis results: (A) *p*-value distribution, (B) log_2_ fold change distribution, (C) relationship between mean expression and log_2_ fold change, and (D) sample size versus statistical significance. Red dashed lines indicate applied significance thresholds.

#### 5.1.4 Hierarchical Clustering of Top DEGs

As shown in Fig. 3, hierarchical clustering based on the top 50 DEGs results in clear segregation between AD and control samples. The observed clustering patterns indicate strong coherence among disease-associated genes.

**Fig. 3:**
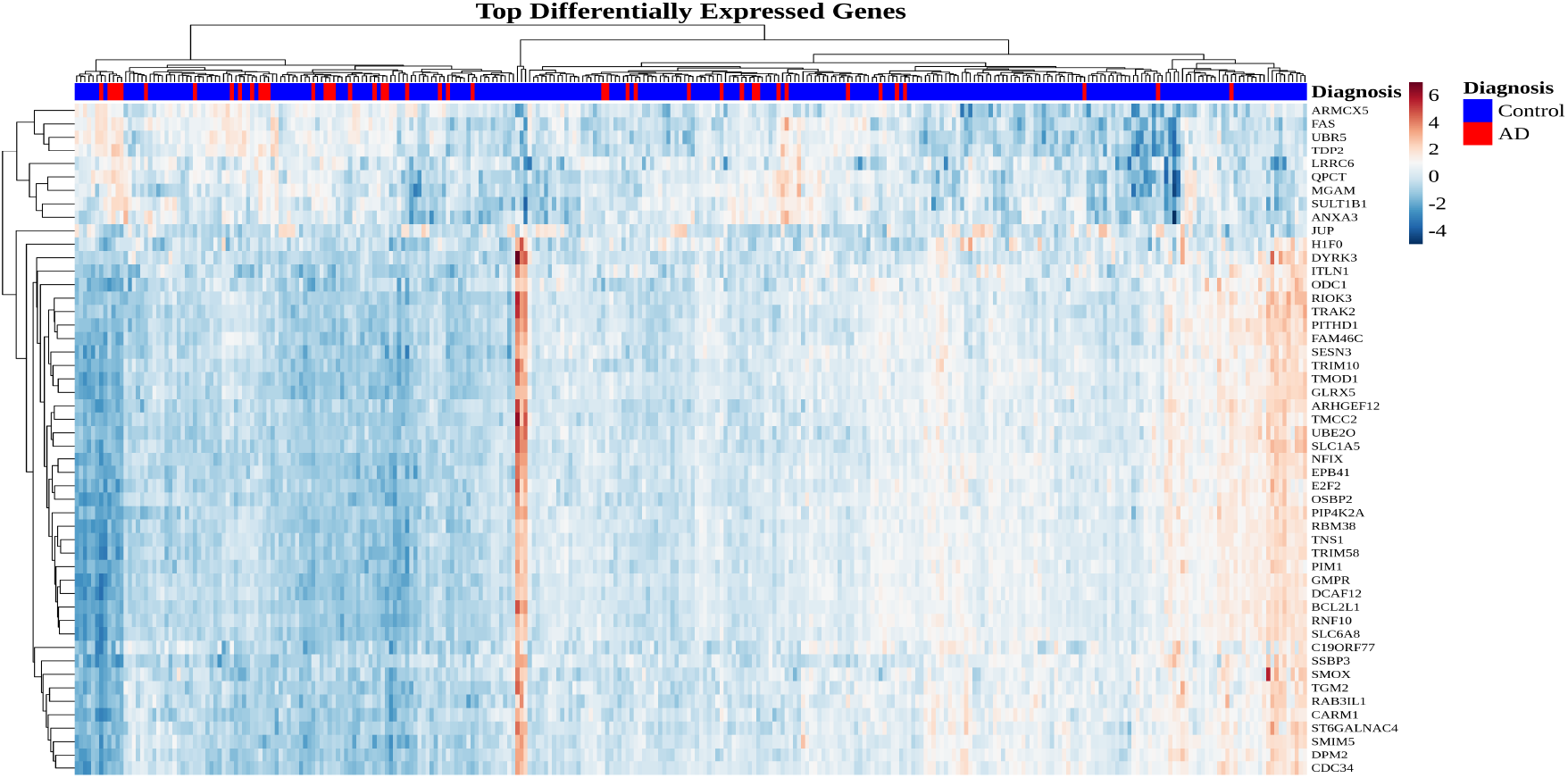
Hierarchical clustering heatmap of the top 50 differentially expressed genes. Rows represent genes and columns represent samples. Expression values are Z-score normalized. The color scale indicates relative expression levels, and dendrograms illustrate hierarchical clustering of both genes and samples.

Overall, the differential expression analysis reveals a robust and asymmetric transcriptional signature associated with AD, characterized predominantly by gene down-regulation. The consistency across statistical distributions, global visualizations, and hierarchical clustering establishes a high-confidence DEG set suitable for downstream functional enrichment and regulatory network analyses.

### 5.2 Target Gene Prediction and miRNA-DEG Overlap Analysis

The objective of this step was to systematically identify DEGs that are potentially regulated by miRNAs, thereby establishing a mechanistic link between post-transcriptional regulation and transcriptional dysregulation observed in AD. This analysis integrates experimentally derived gene expression data from the ADNI cohort with curated miRNA–target interaction datasets to generate a high-confidence set of miRNA–mRNA regulatory relationships.

#### 5.2.1 Overlap Analysis Between DEGs and miRNA Targets

The gene list, when intersected with a comprehensive miRNA target dataset comprising 16,979 genes, the overlap analysis revealed that 15,545 of the 20,092 genes (77.37%) are predicted or experimentally validated targets of one or more miRNAs. This substantial overlap highlights the dominant role of miRNA-mediated post-transcriptional regulation in shaping the AD-associated transcriptomic profile. Out of the 15,545 overlapped genes, a total of 69 downregulated and 22 upregulated genes were targeted, giving us a 61.49% overlap with the significant genes.

A vectorized merging strategy was employed to efficiently construct a detailed miRNA–DEG interaction dataset, resulting in 1,669,089 individual miRNA–target interactions involving 3,055 unique miRNAs. Each interaction was annotated with gene-level expression metrics, regulatory direction, statistical significance, and supporting evidence where available.

#### 5.2.2 Regulatory Direction and Interaction Characteristics

Stratification of miRNA–DEG interactions by regulation direction revealed a pronounced bias toward down-regulated genes.

Although only 69 down-regulated genes were present among the significant DEGs, these genes collectively participated in 11,008 miRNA interactions, indicating dense regulatory targeting. In contrast, the majority of interactions involved genes classified as not statistically significant at the imposed thresholds, reflecting the pervasive baseline activity of miRNAs across the transcriptome.

Highly connected genes such as *DYRK3* emerged as regulatory hubs, being targeted by numerous miRNAs including hsa-miR-26b-5p, hsa-miR-34a-5p, and hsa-miR-27a-3p. These interactions were associated with strong statistical significance and consistent down-regulation, suggesting coordinated miRNA-driven repression.

Table 2 provides a consolidated overview of the target gene prediction and overlap analysis results.

**Table 2:** Summary of Target Gene Prediction and miRNA–DEG Overlap Analysis.

| Analysis Parameter | Value |
| --- | --- |
| Total probes in ADNI dataset | 49,386 |
| Total samples | 748 |
| Valid probe-to-gene mappings | 48,157 (97.5%) |
| Unique DEG-associated gene symbols | 20,092 |
| miRNA target genes analyzed | 16,979 |
| Overlapping DEGs targeted by miRNAs | 15,545 |
| Percentage overlap | 77.37% |
| Total miRNA–DEG interactions | 1,669,089 |
| Unique miRNAs involved | 3,055 |
| Down-regulated genes targeted | 69 |
| Up-regulated genes targeted | 22 |

Collectively, these results demonstrate that a substantial proportion of transcriptional alterations observed in AD are potentially governed by miRNA-mediated regulation. The high density of miRNA–DEG interactions, coupled with the identification of key regulatory hubs, provides a robust foundation for subsequent functional enrichment analysis and network-level modeling.

### 5.3 Functional Enrichment Analysis (FEA)

To elucidate the biological relevance of the miRNA-targeted DEGs identified in Section 5.2, a comprehensive FEA was conducted. The primary objective of this analysis was to identify biological processes, molecular functions, cellular compartments, and signaling pathways that are statistically overrepresented among genes under miRNA-mediated regulatory control in Alzheimer’s disease (AD).

#### 5.3.1 Biological Process Enrichment

Analysis of Gene Ontology Biological Process (GO:BP) terms revealed a strong enrichment of processes related to neuronal communication, synaptic organization, and signal transduction. Prominent biological themes included synaptic signaling, regulation of synaptic plasticity, neurotransmitter secretion, and axon guidance. These processes are central to learning and memory and are known to be severely compromised in Alzheimer’s disease.

In addition to synaptic processes, enrichment was observed for pathways involved in neuronal development, vesicle-mediated transport, and intracellular protein trafficking. The presence of these processes suggests that miRNA-targeted DEGs contribute to the maintenance of synaptic architecture and efficient intracellular communication, both of which are disrupted during neurodegeneration.

Several enriched biological processes were directly associated with cellular stress responses, including regulation of apoptotic signaling, cellular response to oxidative stress, and inflammatory signaling pathways. These findings align with well-established AD pathophysiological mechanisms [36], including progressive neuronal loss, mitochondrial dysfunction, and chronic neuroinflammation.

#### 5.3.2 Molecular Function and Cellular Component Enrichment

Gene Ontology Molecular Function (GO:MF) enrichment analysis demonstrated over-representation of genes involved in protein binding, RNA binding, kinase activity, and transcriptional regulation. These functions indicate that miRNA-regulated genes play key roles in modulating intracellular signaling cascades and gene expression programs, amplifying the downstream effects of miRNA dysregulation [37].

Cellular Component (GO:CC) enrichment analysis revealed that miRNA-targeted DEGs are preferentially localized to neuronal and synaptic compartments. Enriched cellular locations included synaptic membranes, postsynaptic densities, neuron projections, dendrites, and intracellular vesicles. The spatial enrichment of these genes within synaptic structures supports the hypothesis [38] that miRNA-mediated regulation critically influences synaptic integrity and neurotransmission efficiency in AD.

#### 5.3.3 Pathway-Level Enrichment

Pathway enrichment analysis identified significant overrepresentation of multiple signaling pathways implicated in neurodegeneration. Key enriched pathways included PI3K–Akt signaling, MAPK signaling, calcium signaling, neurotrophin signaling, and apoptosis-related pathways. These pathways regulate neuronal survival, synaptic plasticity, and responses to cellular stress, all of which are perturbed in AD [39].

Notably, pathways associated with vesicle trafficking, endocytosis, and protein processing were also enriched, suggesting that miRNA regulation may influence amyloid precursor protein (APP) trafficking and processing. This observation provides a mechanistic link between miRNA dysregulation and amyloidogenic processes central to AD pathology [40].

#### 5.3.4 Biological Implications

The functional enrichment results demonstrate that miRNA-targeted DEGs are not randomly distributed across functional categories but instead converge on biologically coherent pathways central to neuronal function and survival. The coordinated enrichment of synaptic, signaling, and apoptotic processes indicates that miRNAs act as higher-order regulators capable of modulating entire molecular programs rather than isolated genes [41].

Importantly, the enrichment of pathways related to synaptic transmission and intracellular signaling provides functional validation for the miRNA–mRNA interactions identified in Section 5.2. These results suggest that miRNA dysregulation contributes to Alzheimer’s disease progression by simultaneously impairing neuronal communication, promoting cellular stress, and weakening neuroprotective signaling networks.

In a nutshell, functional enrichment analysis revealed that miRNA-targeted DEGs are significantly enriched in neuronal, synaptic, and signaling-related biological processes, as well as pathways governing apoptosis and cellular stress responses. These findings establish a strong functional context for the identified miRNA–DEG interactions and provide a mechanistic foundation for the regulatory network construction described in the subsequent section.

### 5.4 Network-Based Centrality and Backbone Analysis

To elucidate the organizational principles and regulatory hierarchy governing the miRNA–gene interaction landscape, a comprehensive network-based analysis was performed. The miRNA-gene interaction network was modeled as a bipartite graph, with miRNAs and protein-coding genes represented as distinct node classes and edges denoting high-confidence regulatory interactions. Multiple centrality metrics were computed to quantify node importance from complementary topological perspectives, followed by backbone extraction to isolate dominant regulatory cores.

#### 5.4.1 Global Filtered miRNA-Gene Network

The filtered miRNA-gene network comprises 2,343 nodes connected by 14,603 edges, including 2,252 miRNAs and 91 significant genes. The resulting topology is sparse (density = 0.0053) yet highly centralized, reflecting a regulatory architecture dominated by a limited set of influential hubs.

#### 5.4.2 Hub miRNA Characterization via Ego Networks

To investigate local regulatory influence, ego networks were constructed for the most highly connected miRNAs. Each ego network consists of a central miRNA node and its directly connected gene targets, enabling direct visualization of first-order regulatory reach. Fig. 4 shows the most prominent ego networks.

**Fig. 4:**
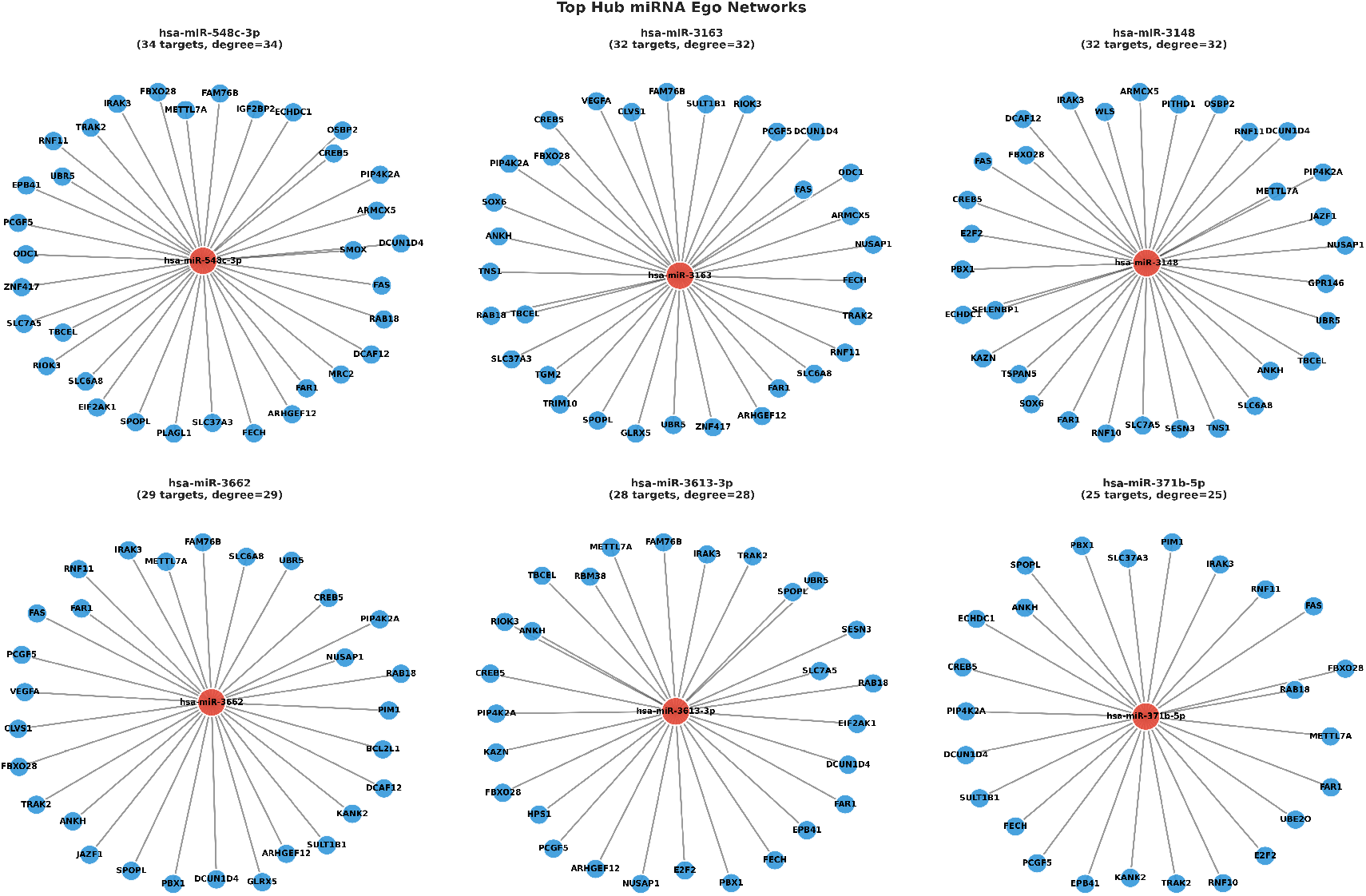
Ego networks of top hub miRNAs ranked by degree centrality. Central nodes represent hub miRNAs, while peripheral nodes denote directly regulated gene targets.

Prominent hub miRNAs, including *hsa-miR-548c-3p, hsa-miR-3163, hsa-miR-3148*, and *hsa-miR-3662*, regulate between 25 and 34 genes each, exhibiting star-like network configurations characteristic of master post-transcriptional regulators.

#### 5.4.3 Centrality Metrics and Hub Ranking

To quantitatively assess node importance, degree, betweenness, closeness, and eigenvector centrality measures were computed. Degree centrality identifies highly connected hubs, betweenness centrality captures nodes acting as inter-module bridges, closeness centrality reflects global reachability, and eigenvector centrality measures influence derived from connections to other highly ranked nodes. Table 3 shows the top central nodes in the network. The dominance of PBX1 and SLC7A5 across degree, betweenness, and eigenvector centrality underscores their dual role as both highly connected hubs and critical conduits of regulatory information. In contrast, closeness centrality is largely dominated by miRNAs, reflecting their ability to rapidly influence large portions of the network.

**Table 3:** Top-ranked nodes across multiple centrality metrics in the miRNA–gene regulatory network.

| Metric | Rank | Node | Node Type | Centrality Value |
| --- | --- | --- | --- | --- |
| Degree | 1 | PBX1 | Gene | 591 |
| Degree | 2 | SLC7A5 | Gene | 527 |
| Degree | 3 | TNS1 | Gene | 393 |
| Degree | 4 | PIP4K2A | Gene | 367 |
| Degree | 5 | PCGF5 | Gene | 364 |
| Degree | 6 | ZNF417 | Gene | 356 |
| Degree | 7 | RNF11 | Gene | 349 |
| Degree | 8 | SPOPL | Gene | 336 |
| Degree | 9 | E2F2 | Gene | 312 |
| Degree | 10 | FBXO28 | Gene | 312 |
| Betweenness | 1 | SLC7A5 | Gene | 0.1069 |
| Betweenness | 2 | PBX1 | Gene | 0.0971 |
| Betweenness | 3 | RNF11 | Gene | 0.0655 |
| Betweenness | 4 | ZNF417 | Gene | 0.0611 |
| Betweenness | 5 | VEGFA | Gene | 0.0597 |
| Closeness | 1 | hsa-miR-1976 | miRNA | 0.4312 |
| Closeness | 2 | hsa-miR-3148 | miRNA | 0.4295 |
| Closeness | 3 | hsa-miR-548c-3p | miRNA | 0.4282 |
| Eigenvector | 1 | PBX1 | Gene | 0.2313 |
| Eigenvector | 2 | SLC7A5 | Gene | 0.1763 |
| Eigenvector | 3 | TNS1 | Gene | 0.1553 |

#### 5.4.4. Integrated Backbone Comparison

Network centrality analysis revealed two complementary regulatory architectures as evident from Fig. 5. The betweenness-based backbone highlighted a small set of miRNAs and genes acting as bridges between distinct regulatory regions, suggesting their potential role in coordinating cross-module communication. In contrast, the degree-based backbone was dominated by classical hub regulators exhibiting extensive connectivity, consistent with their role as major transcriptional or post-transcriptional controllers.

**Fig. 5:**
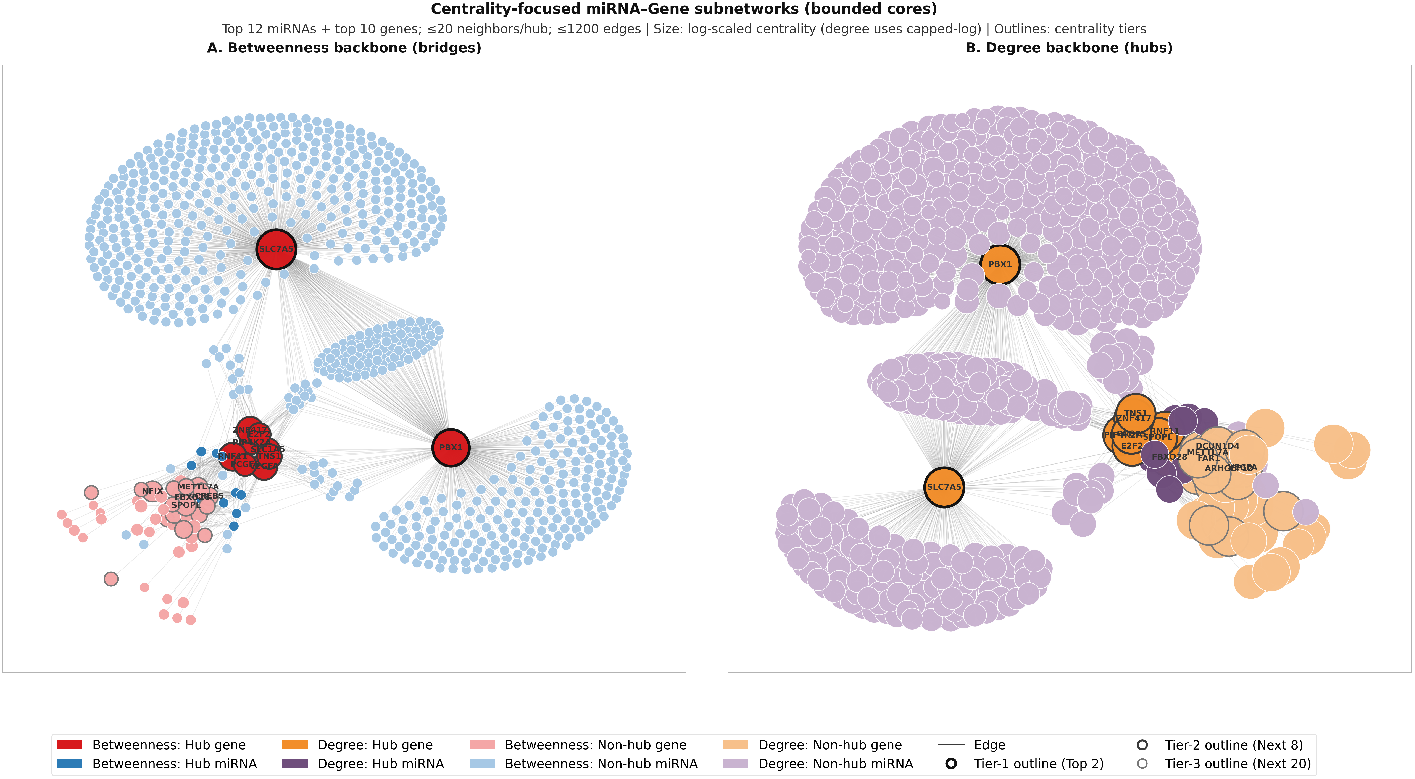
Centrality-driven miRNA–gene regulatory backbones. The miRNA–gene regulatory network was reduced to centrality-focused backbone subnetworks to highlight key regulatory structures while avoiding visual overcrowding.(A) Betweenness backbone: Nodes with high betweenness centrality were selected to identify bridge regulators that connect otherwise weakly linked gene–miRNA modules. These nodes represent potential bottlenecks or mediators of information flow in the regulatory network. (B) Degree backbone: Nodes with high degree centrality were selected to identify hub regulators with extensive connectivity, representing highly influential miRNAs or genes that regulate many targets. In both panels, node size is proportional to the corresponding centrality measure, blue nodes represent genes, grey nodes represent miRNAs, and red outlines indicate top-ranked central regulators. To ensure interpretability, each backbone was constructed using a bounded neighborhood strategy (≤20 neighbors per hub, ≤1200 edges).

Notably, several nodes appear in both backbones, indicating regulators that are not only highly connected but also critical for network-wide information flow. These nodes may represent particularly important candidates for functional validation or therapeutic targeting.

### 5.5 Machine Learning Classification Performance

The performance of the proposed machine learning framework was evaluated using standard classification metrics, including accuracy, ROC–AUC, precision, sensitivity (recall), specificity, and F1-score. Model evaluation was performed using stratified five-fold cross-validation on the training data, followed by independent test-set evaluation to assess generalization capability for distinguishing Alzheimer’s disease (AD) samples from control samples. Table 4 shows the five-fold cross-validation training performance of machine learning models.

**Table 4:**
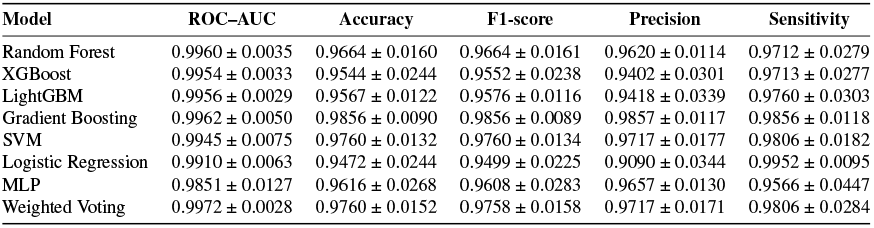
Five-fold cross-validation (training) performance of machine learning models (mean ± standard deviation)

| Model | ROC-AUC | Accuracy | F1-score | Precision | Sensitivity |
| --- | --- | --- | --- | --- | --- |
| Random Forest | 0.9960 ± 0.0035 | 0.9664 ± 0.0160 | 0.9664 ± 0.0161 | 0.9620 ± 0.0114 | 0.9712 ± 0.0279 |
| XGBoost | 0.9954 ± 0.0033 | 0.9544 ± 0.0244 | 0.9552 ± 0.0238 | 0.9402 ± 0.0301 | 0.9713 ± 0.0277 |
| LightGBM | 0.9956 ± 0.0029 | 0.9567 ± 0.0122 | 0.9576 ± 0.0116 | 0.9418 ± 0.0339 | 0.9760 ± 0.0303 |
| Gradient Boosting | 0.9962 ± 0.0050 | 0.9856 ± 0.0090 | 0.9856 ± 0.0089 | 0.9857 ± 0.0117 | 0.9856 ± 0.0118 |
| SVM | 0.9945 ± 0.0075 | 0.9760 ± 0.0132 | 0.9760 ± 0.0134 | 0.9717 ± 0.0177 | 0.9806 ± 0.0182 |
| Logistic Regression | 0.9910 ± 0.0063 | 0.9472 ± 0.0244 | 0.9499 ± 0.0225 | 0.9090 ± 0.0344 | 0.9952 ± 0.0095 |
| MLP | 0.9851 ± 0.0127 | 0.9616 ± 0.0268 | 0.9608 ± 0.0283 | 0.9657 ± 0.0130 | 0.9566 ± 0.0447 |
| Weighted Voting | 0.9972 ± 0.0028 | 0.9760 ± 0.0152 | 0.9758 ± 0.0158 | 0.9717 ± 0.0171 | 0.9806 ± 0.0284 |

Table 5 shows the five-fold cross-validation training performance of machine learning models. The corresponding heatmap is shown in Fig. 6. Fig. 7 shows the ROC-AUC of the models on the independent test set. The visualization highlights the comparative strengths of ensemble and boosted models, with XGBoost and LightGBM achieving consistently high accuracy, specificity, and F1-scores, while logistic regression demonstrates superior ROC–AUC performance.

**Table 5:** Independent test-set performance of machine learning models for AD classification.

| Model | Threshold | Accuracy | ROC-AUC | Precision | Sensitivity | Specificity | F1-score |
| --- | --- | --- | --- | --- | --- | --- | --- |
| Random Forest | 0.4935 | 0.9344 | 0.9017 | 0.7778 | 0.7778 | 0.9615 | 0.7778 |
| XGBoost | 0.4964 | 0.9508 | 0.9295 | 0.8750 | 0.7778 | 0.9808 | 0.8235 |
| LightGBM | 0.5566 | 0.9508 | 0.9081 | 0.8750 | 0.7778 | 0.9808 | 0.8235 |
| Gradient Boosting | 0.0586 | 0.9180 | 0.9103 | 0.7000 | 0.7778 | 0.9423 | 0.7368 |
| SVM | 0.0117 | 0.8689 | 0.9359 | 0.5333 | 0.8889 | 0.8654 | 0.6667 |
| Logistic Regression | 0.9329 | 0.9344 | 0.9594 | 0.7778 | 0.7778 | 0.9615 | 0.7778 |
| MLP | 0.9776 | 0.9180 | 0.8889 | 0.8333 | 0.5556 | 0.9808 | 0.6667 |
| Weighted Voting | 0.3244 | 0.9180 | 0.9017 | 0.7000 | 0.7778 | 0.9423 | 0.7368 |

**Fig. 6:**
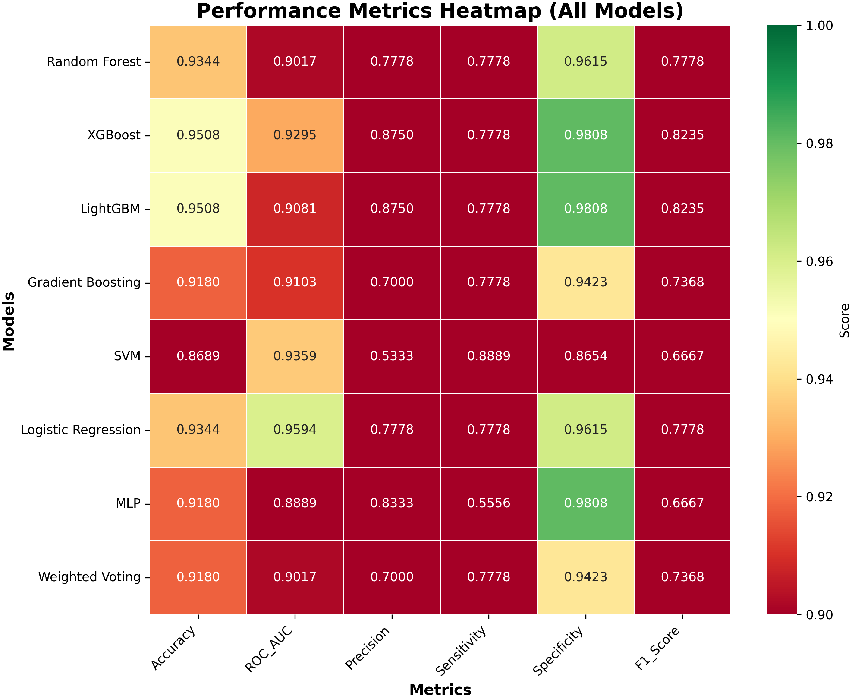
Heatmap visualization of key performance metrics, including accuracy, ROC–AUC, precision, sensitivity, specificity, and F1-score, for all evaluated machine learning models on the test set.

**Fig. 7:**
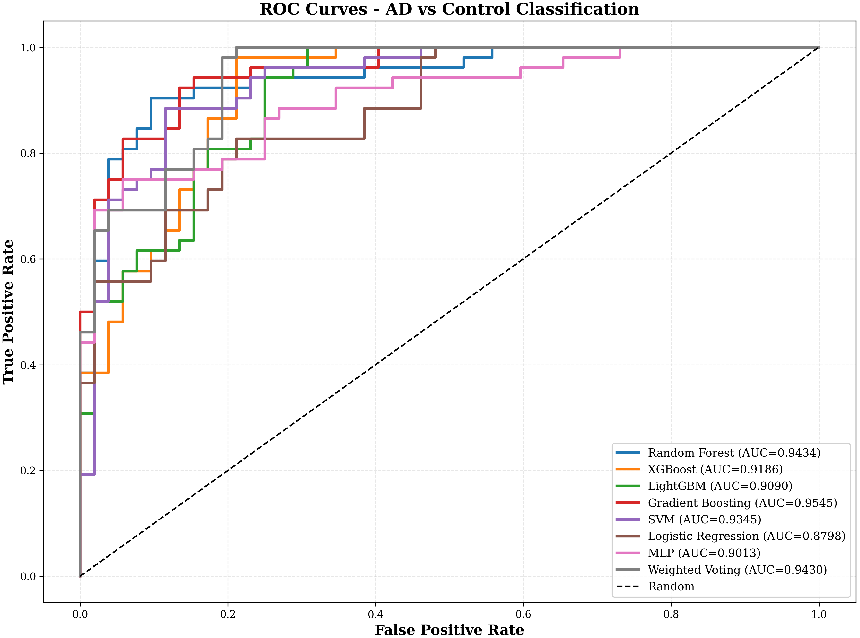
Receiver Operating Characteristic (ROC) curves for all machine learning models evaluated on the test set for Alzheimer’s disease (AD) versus control classification. The area under the ROC curve (AUC) values indicates strong discriminative capability across models, with logistic regression and XGBoost achieving the highest AUC scores, confirming their effectiveness in separating AD and control samples across varying decision thresholds.

The confusion metrics of the models on the test set are shown in Fig. 9. Ensemble and gradient-based models, particularly XGBoost and LightGBM, exhibit lower misclassification rates compared to other classifiers, highlighting their robustness in distinguishing AD from control samples. The classification report for the best model, XGBoost, is shown in Fig. 8. The model demonstrates high precision and recall for the Control class, while maintaining strong discriminative performance for AD samples, indicating a balanced and reliable classification behavior.

**Fig. 8:**
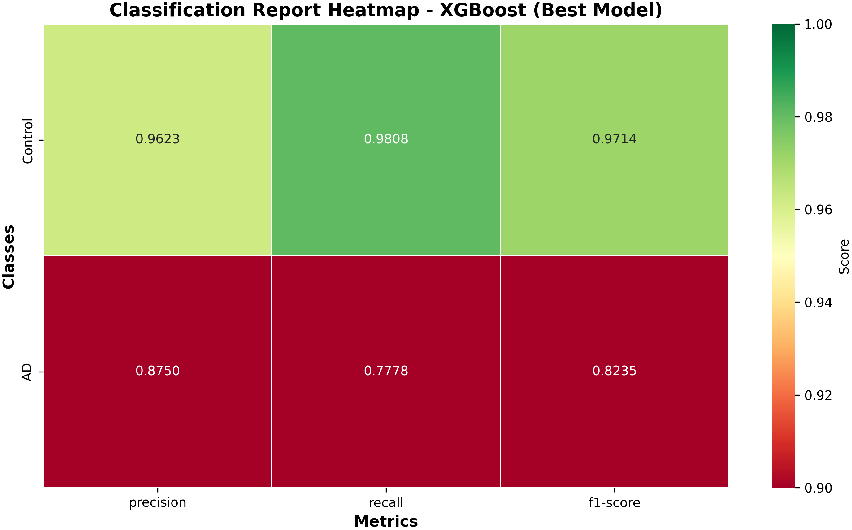
Classification report heatmap for the best-performing model (XGBoost) on the independent test set, illustrating class-wise precision, recall, and F1-score for Control and Alzheimer’s disease (AD) samples.

**Fig. 9:**
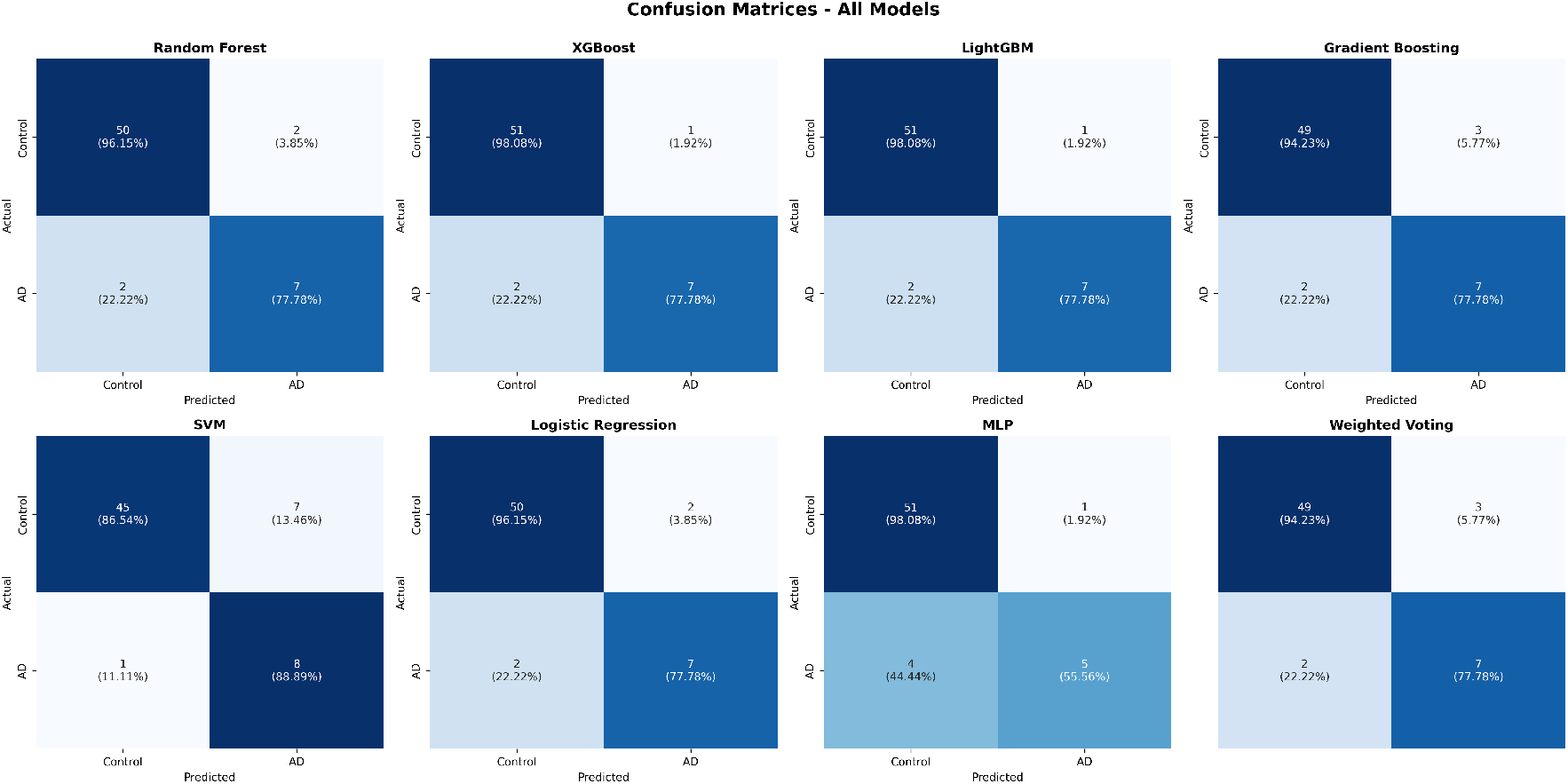
Confusion matrices for all evaluated machine learning models on the test set, showing the distribution of true positives, true negatives, false positives, and false negatives for Control and Alzheimer’s disease (AD) classes.

Overall, the cross-validation results demonstrate consistently high ROC–AUC, accuracy, and sensitivity across all models, indicating strong robustness and minimal variance during training. Independent test-set evaluation further confirms the generalization capability of ensemble and gradient-based models, with XGBoost and LightGBM achieving the highest accuracy and F1-scores, while logistic regression yielded the highest ROC–AUC. These findings highlight the effectiveness of ensemble learning and boosted decision-tree models for reliable Alzheimer’s disease classification.

## 6 Conclusion

This study presents an integrated computational framework that combines differential expression analysis, miRNA–mRNA regulatory network modeling, functional enrichment analysis, and supervised machine learning to investigate AD-associated molecular mechanisms. A relevant set of DEGs was identified and intersected with the high-confidence miRNA targets, identifying the disease-relevant regulatory interactions and hub regulators with prominent network centrality. FEA indicated that these targets converge on synaptic signaling, intracellular transport, stress-response, and neurodegeneration-related pathways, thereby supporting their biological relevance to AD pathology.

ML models trained on the selected molecular features achieved consistent classification performance, as reflected by the various classification report metrics (precision, recall, F1-score, and accuracy) and the confusion matrix, which indicated the effective separation between AD and control samples. The ROC-AUC curve further highlighted the discriminative power of these models by pointing out the existence of a balance between sensitivity and specificity. Evaluation on the independent test set demonstrates the robust generalization of the ensemble and gradient-based models, with XGBoost and LightGBM showing the top accuracy and F1-scores, and logistic regression exhibiting the highest ROC–AUC. Taken together, these results establish the fact that the concept of integrative network-guided transcriptomics and ML models has the capability of generating molecular signatures that are biologically interpretable with potential utility for AD characterization and biomarker discovery.

## Acknowledgement

Data used in preparation of this article were obtained from the Alzheimer’s Disease Neuroimaging Initiative (ADNI) database (adni.loni.usc.edu). As such, the investigators within the ADNI contributed to the design and implementation of ADNI and/or provided data but did not participate in the analysis or writing of this report. A complete listing of ADNI investigators can be found at: http://adni.loni.usc.edu/wp-content/uploads/how_to_apply/ADNI_Acknowledgement_List.pdf

